# Logical and experimental modeling of cytokine and eicosanoid signaling in psoriatic keratinocytes

**DOI:** 10.1101/2021.06.07.447313

**Authors:** Eirini Tsirvouli, Felicity Ashcroft, Berit Johansen, Martin Kuiper

## Abstract

Psoriasis is characterized by chronic inflammation, perpetuated by a Th17-dependent signaling loop between the immune system and keratinocytes that could involve phospholipase A_2_ (PLA_2_)-dependent eicosanoid release. A prior knowledge network supported by experimental observations was used to encode the regulatory network of psoriatic keratinocytes in a computational model for studying the mode of action of a cytosolic (c) PLA_2_α inhibitor. A combination of evidence derived from the computational model and experimental data suggests that Th17 cytokines stimulate pro-inflammatory cytokine expression in psoriatic keratinocytes via activation of cPLA_2_α-PGE_2_-EP4 signaling, which could be suppressed using the anti-psoriatic calcipotriol. cPLA_2_α inhibition and calcipotriol showed overlapping and distinct modes of action. Model analyses revealed the immunomodulatory role of Th1 cytokines, the modulation of the physiological states of keratinocytes by Th17 cytokines, and how Th1 and Th17 cells together promote the development of psoriasis. Model simulations additionally suggest novel drug targets, including EP4 and PRKACA, for treatment that may restore a normal phenotype. Our work illustrates how the study of complex diseases can benefit from an integrated systems approach.

## 1. Introduction

Psoriasis is a chronic inflammatory disease that affects 2-3% of the world’s population. Psoriasis vulgaris, commonly referred to as plaque-type psoriasis, is the most prominent type of the disease accounting for 90% of total cases (Boehncke & Schön, 2015). The characteristic histological changes of the disease are the hyperproliferation and aberrant differentiation of keratinocytes (KCs), together with the inflammatory state and infiltration of immune cells in the dermis and epidermis (Meephansan *et al*, 2017). While the infiltration of immune cell subpopulations, namely dendritic cells (DCs), T-lymphocytes, neutrophils and mast cells, were for a long time considered the main pathogenic factor of psoriasis, the key role of KCs in the initiation and the maintenance of the chronic nature of the disease is indisputable (Lowes *et al*, 2013). Therefore, the disease appears to be a result of a rather complex interaction between different cell types (Georgescu *et al*, 2019; Benhadou *et al*, 2019; Albanesi *et al*, 2018; Deng *et al*, 2016) and should be studied as a multiscale dysregulation. The understanding of the complex dynamics of psoriasis, and the identification of effective treatments may benefit from an integrative approach and the construction of systems-based models.

Psoriasis can be divided into two main stages: initiation and maintenance/chronic stage. In early events, KCs respond to different environmental and genetic stimuli by secreting immune mediators that result in the recruitment of DCs to the dermis and epidermis and their subsequent maturation. Activated DCs are then able to secrete cytokines that influence the differentiation of naive T-cells (Albanesi *et al*, 2018). In psoriasis, three T-helper (Th) cell subpopulations have well-documented roles in the progression and maintenance of the disease: Th1, Th17, and Th22 cells. Each Th type contributes to the cytokine milieu of psoriatic lesions. Th1 cells release IFNγ and TNFα, while Th17 produce IL-17 and IL-22, partially overlapping with the role of Th22 cells that produce only IL-22 (Benham *et al*, 2013, 22). The activation of these three classes of immune cells, in turn, triggers KCs to further produce antimicrobial peptides (AMPs), proinflammatory cytokines, and chemoattractants, which all together recruit neutrophils and maintain the same Th-cell subpopulation in a positive feedback loop that sustains chronic inflammation (Georgescu *et al*, 2019; Albanesi *et al*, 2018; Deng *et al*, 2016).

The chronic, life-long nature of psoriasis requires long-term treatment plans that are able to cope with the relapsing episodes of the disease (Gisondi *et al*, 2017). The latest advances in our understanding of the disease have led to the development of biologics targeting specific immune components (cells or proteins) (Kamata & Tada, 2020). Other treatment options include vitamin D analogues such as calcipotriol, which affects KCs and immune cells (Kim, 2010). Current treatment options appear to have relatively high efficacy and tolerability, however, immediate or long-lasting efficacy can vary between patients (Kamata & Tada, 2020; Kim, 2010), and certain biologics are accompanied by side effects and tolerability issues such as recurrent infections (Kamata & Tada, 2020). Combination treatment with two or more agents has been proposed as a way to overcome these challenges and improve efficacy in non-responding patients while ameliorating and reducing the potential side effects of long-term treatments (Cather & Crowley, 2014).

The dysregulated hydrolysis of phospholipids catalyzed by the phospholipase A2 (PLA2) enzyme family is involved in multiple diseases, including cancer, rheumatoid arthritis, and psoriasis (Magrioti & Kokotos, 2013; Leistad *et al*, 2011; Linkous & Yazlovitskaya, 2010). Cytosolic PLA2*a* (cPLA_2_α) hydrolyzes phosphatidylcholine (PC) to yield lysophosphatidylcholine (LPC) and arachidonic acid (AA), two bioactive lipids, the latter of which can be further metabolized by cyclooxygenase (COX) and lipoxygenase (LOX) enzymes to produce prostaglandins (PGs), leukotrienes (LTs) and hydroxy fatty acids e.g. 12s-HETE. These AA metabolites, or eicosanoids, are involved in various diseases, often producing inflammatory responses, but can also be involved in the activation of survival-promoting kinases (Linkous & Yazlovitskaya, 2010; Hirabayashi *et al*, 2004; Murakami *et al*, 2017) and cell cycle progression (Naini *et al*, 2015, 2016).

The signaling through cPLA_2_*a* and its downstream lipid mediators has a prominent role in different stages of psoriasis, with epidermal KCs being an important source of eicosanoids, including prostanoids and leukotrienes (Nicolaou, 2013), which similarly participate in the infiltration and amplification of certain immune cell types (Ueharaguchi 2018, Camp et al 1984; Sheibanie 2004; Lee, 2019) and act upon the KCs themselves via multiple prostanoid and leukotriene receptors (Konger 1998, Konger 2005; Kanda 2007, Kim 2010.) While Th1 cytokines are established activators of phospholipases and cause eicosanoid release from KCs (Sjursen *et al*, 2000; Thommesen *et al*, 1998) the effects of Th17 cytokines on eicosanoid production and signaling is unclear. Furthermore, PGE_2_ has been reported to regulate Th17 and Th1 differentiation and response (Boniface *et al*, 2009; Napolitani *et al*, 2009; Yao *et al*, 2009), as well as affecting DCs (Chizzolini & Brembilla, 2009). PGE_2_ acts through four receptors, EP1-EP4, all expressed in KCs (Rundhaug *et al*, 2011). The main receptor involved in psoriasis is EP4 (Tsuge *et al*), which activates both the AC/cAMP pathway and the PI3K/AKT pathway (Rundhaug *et al*, 2011). While the roles of EP1 and EP3 in psoriasis have not been well studied, their associated phenotypes do not appear to be characteristic for the disease as EP1 was found to promote differentiation in non-melanoma KCs (Konger *et al*, 2009) and EP3 was proposed as a growth inhibitor for KCs (Konger *et al*, 2005).

The targeting of cPLA_2_*a* with chemical inhibitors has been suggested as a promising treatment against hyperproliferative and inflammatory diseases (Yarla *et al*, 2016), also supported by the promising results in clinical trials against mild to moderate psoriasis (Omland *et al*, 2017). However, the limited understanding of the exact involvement of cPLA_2_*a* in psoriasis, as well as the general gaps of knowledge of its involvement in the development of the disease, require a unified framework that collects prior knowledge and integrates new observations to produce a modeling tool that can be further used to not only make sense of observations but also rationalize experimental decisions, test hypotheses and highlight gaps of knowledge. For this reason, *in silico* approaches to predict and interpret the effects of various stimuli and perturbations in cellular systems have become increasingly relevant. Among different possible approaches, logical models that represent the signaling events and causal interactions between cellular components prove to be able to capture the behavior of cells upon various experiments, predict the effect of drugs, and identify potentially synergistic combinations. (Tsirvouli *et al*, 2020; Pirkl *et al*, 2016; Flobak *et al*, 2015).

In this study, we aimed to develop such a logical model to represent the complex signaling events occurring in psoriatic keratinocytes to (1) investigate the mode of action of a cPLA_2_*a* inhibitor (2) predict therapeutic benefits of drug combinations, and (3) investigate the contributions of Th17 versus Th1 cytokines to disease phenotypes.

## 2. Methods

### Section I: *In vitro* methods

#### 2.1.1 Materials

The reagents used for culturing cells, and other chemicals, including calcipotriol hydrate were purchased from Sigma-Aldrich (St. Louis, MPS, US) unless stated otherwise. The fluoroketone AVX001 was synthesized and characterized according to Holmeide and Skattebol and provided by Dr. Inger Reidun Aukrust and Dr. Marcel Sandberg (Synthetica AS, Norway). AVX001 was stored at −80 °C as a 20 mM stock solution in DMSO under argon gas to minimize oxidation. Interleukins 17A (#8928SC) and 22 (#8931SC) were purchased from Cell Signalling Technologies. Recombinant human epidermal growth factor (EGF) was from R&D Systems (Abingdon, UK).

#### 2.1.2 Maintenance of HaCaT Keratinocytes

The spontaneously immortalized skin KC cell line HaCaT (Boukamp *et al*, 1988) was kindly provided by Prof. N. Fusenig (Heidelberg, Deutsches Krebsforschungszentrum, Germany). The cells were maintained in Dulbecco’s modified Eagle Medium (DMEM) supplemented with 5% (v/v) FBS, 0.3 mg/mL glutamine, and 0.1 mg/mL gentamicin (DMEM-5) at 37 °C with 5% CO2 in a humidified atmosphere at sub-confluency. HaCaT are commonly used cell system to study proliferative and inflammatory responses in psoriasis research (Farkas *et al*, 2003; Feuerherm *et al*, 2013; Lehmann, 1997; Pozzi *et al*, 2004; Se *et al*, 2010; Ziv *et al*, 2008); the cells differentiate on reaching confluency and form stratified epithelia when grown under appropriate conditions (Hinitt *et al*, 2011; Schoop *et al*, 1999)

#### 2.1.3 3D Culture of HaCaT Keratinocytes

3D culture of HaCat KCs was carried out using the 24-well carrier plates system from Thermo Fisher Scientific (#141002) according to the protocol described (Ashcroft *et al*, 2020). Briefly, the culture inserts (0.4 μm pore size) were coated using the Coating Matrix kit (Thermofisher Scientific #R-011-K) according to the manufacturer’s protocol. HaCaT were seeded at a density of 0.3 × 10^5^ cells per insert in 0.5 mL DMEM-5. and incubated for 24 h before being lifted to the air-liquid interface. The media in the lower chamber was replaced with DMEM-5 (without antibiotics) + 1 ng/mL EGF, and 5 μg/mL L-ascorbic acid in the absence or presence of Th17 psoriatic cytokines (IL-17, 10 ng/ml and IL-22, 10 ng/ml) and either AVX001 (5 μM), calcipotriol (10 nM) or a combination of AVX001 (5 μM) and calcipotriol (10 nM). The media in the lower chambers was changed every 3rd day for 12 days.

#### 2.1.4 RNA Extraction and Real-Time QuantitativePCR

Total RNA was extracted from HaCaT cultures using the RNeasy kit (QIAGEN) according to the manufacturer’s protocol. The amount and purity of the RNA samples were quantified using a Nanodrop One/OneC Microvolume UV–VIS Spectrophotometer (ND-ONE-W) from ThermoFisher Scientific. RNA samples with absorbance (A) 260/280 between 2.0 and 2.2 were accepted. Reverse transcription was carried out using the QuantiTect Reverse Transcription Kit (Qiagen, Hilden, Germany) according to the manufacturer’s protocol, using 1 μg of total RNA per sample. Real-time PCR analysis was performed using the LightCycler 480 SYBR Green I Master MIX and LightCycler 96 instrument from Roche (Basel, Switzerland), according to the manufacturer’s protocol with a final individual primer concentration of 0.5 μM. Amplification was performed with an initial step of 95 °C (10 min) followed by 45 cycles of denaturation at 95 °C (10 s), annealing at 55°C (10 s) and elongation at 72 °C (10 s). Melting analysis was performed with the following parameters: 95 °C for 5 s, 65 °C for 1 min, 97 °C for 1 min and cooling at 50 °C for 10 s. Three reference genes were selected for gene expression analysis: TBP, HPRT1, and GAPDH (Allen *et al*, 2008).) All primers were KiCqStart primers designed by and purchased from Sigma Aldrich. Ct values and PCR amplification efficiencies were calculated using LinRegPCR software (version 2017.1) (Ruijter *et al*, 2014; Tuomi *et al*, 2010) and imported into qBASE+ (version 3.2) (Hellemans *et al*, 2007) for relative expression analysis.

#### 2.1.5 Immunohistochemistry for analysis of proliferation and differentiation markers

HaCaT cultures were processed in parallel to quantify the expression of ki67 and cytokeratin 10 as determinants of the proportion of proliferating cells and cells undergoing differentiation. These cultures were fixed by immersion in 4% paraformaldehyde (PFA) overnight; the membranes were removed from the inserts and prepared in Tissue Clear (Sakura, Osaka, Japan) for paraffin wax embedding using the Excelsior AS Tissue processor (ThermoFisher Scientific). Paraffin-embedded sections (4 μm) were cut onto SuperFrost Plus slides (ThermoFisher scientific), dried overnight at 37 °C, and then baked for 60 min at 60 °C. The sections were dewaxed in Tissue Clear and rehydrated through graded alcohols to water in an automatic slide stainer (Tissue-Tek Prisma, Sakura). Next, the sections were pretreated in Target Retrieval Solution, High pH (Dako, Glostrup, Denmark, K8004) in PT Link (Dako) for 20 min at 97 °C to facilitate antigen retrieval. The staining was performed according to the manufacturer’s procedure with EnVision G|2 Doublestain System Rabbit/Mouse (DAB+/Permanent Red) kit (Dako/Agilent K5361) on the Dako Autostainer. Endogenous peroxidase and alkaline phosphatase activity were quenched with Dual Endogenous Enzyme Block (Dako). Sections were then rinsed in wash buffer and incubated with primary antibody against Ki67 (MIB1 (Dako M7240) diluted 1:300) for 40 min. The slides were rinsed before incubating in horseradish peroxidase (HRP) - polymer and 3,3’-Diaminobenzidine (DAB) to develop the stain. After a double stain block, the sections were incubated in antibody against cytokeratin 10 (Invitrogen #MA5-13705 diluted 1:100) for 60 min. After incubation in the mouse/rabbit linker, the sections were incubated in AP- polymer and the corresponding red substrate buffer with washing between each step. Tris-buffered saline (TBS; Dako K8007) was used throughout the washing steps. The slides were lightly counterstained with hematoxylin, completely dried, and coverslipped. Appropriate negative controls were performed; both mouse monoclonal isotype control (Biolegend, San Diego, CA) and omitting the primary antibody (negative method control). Images were taken at 40X magnification and >20 images were typically collected per sample for counting.

#### 2.1.6 Enzyme-Linked Immunoassay Detection of Eicosanoids

Supernatants from the HaCaT cultures were analyzed by enzyme-linked immunosorbent assay (ELISA) for PGE_2_ (Cayman #514435), LTB_4_ (Cayman #10009292), TxB_2_ (Cayman #501020), and 12s-HETE (Enzo Life Sciences #ADI-900-050) according to the manufacturer’s protocols. Cell supernatants were assayed at dilutions of 1:100 for PGE_2_, except supernatants from non-LPS-treated PBMC that were assayed undiluted in all assays. Supernatants were hybridized overnight, and the enzymatic conversion of the substrate was read at OD420 nm. Data were processed using a 4-parameter logistic fit model.

#### 2.1.7 Experimental data discretization

In order to make the expression level measured by qPCR comparable to the discrete (i.e. 0 and 1) states that the model predicts, a discretization step was performed. As there is no specific threshold for inferring the state of the different gene products, the state was based on the relative expression of the genes. The discretization workflow is presented in Figure S1 in the Supplementary Material.

### Section II: *In silico* part

#### 2.2.1 Representation of signaling network in psoriatic KCs

A Prior Knowledge Network (PKN) encompassing the main signaling events taking place in KCs during the chronic stage of psoriasis was manually curated based on prior knowledge of the disease. The relevant to the system biological entities (*nodes*) and their interactions (*edges*) were identified by an extensive and careful assessment of the available literature and by their extraction from databases with causal molecular interactions, such as Signor (Perfetto et al. 2016). The metabolism of those eicosanoids with a reported role in psoriasis and those identified through our in vitro experiments was also added to the keratinocyte model. The rest of the curation was focused on the signaling cascades initiated by the main cytokines that characterize psoriatic lesions in the chronic stage of the disease, namely IL-17, IL-22, TNFα, and IFNγ. Additionally, guided by *in vitro* observations (presented in Section 3.1), signaling mechanisms that occur downstream of PGE_2_ were added to the model to allow for hypothesis-driven simulations. In order to allow the exploration of the potential synergistic effect between cPLA_2_*a* inhibition and vitamin D analogs like calcipotriol, signaling through the Vitamin-D receptor (VDR) was included. Each signaling component was annotated by its official gene symbol, UniProt ID or ChEBI ID, while all interactions were annotated with the PubMed IDs supporting their addition to the model.

#### 2.2.2 The logical model of psoriatic keratinocytes

After the identification of the relevant nodes and their regulatory interactions with other nodes, the network was converted to a Boolean model and encoded in the GINsim software (Naldi *et al*, 2018). In Boolean models, the activity of a node (also called *state*) can be associated with Boolean values (i.e. 1 or 0). Signaling components with a value of 1 are considered active while a value of 0 is associated with inactive components. Lastly, for each node, the logical rules governing its state were specified. The logical rules aimed to capture how a node’s state depends on the state and combinations of its regulators and were manually specified following the logical formalism AND, OR, and NOT. The definition of the rules was an iterative process based on the results of various model analyses and their comparison to the expected behavior described in the literature and in our *in vitro* observations.

The final model was analyzed using software tools developed by the CoLoMoTo consortium (http://www.colomoto.org/). The analyses explored the system from several angles aiming i) to assess its performance and validate its closeness to biological reality, ii) to use the model as a tool to understand how cPLA_2_*a* downstream lipid mediators and more specifically PGE_2_, together with Th17 and Th1 cytokines contribute to the different phenotypes of the disease, and iii) to explore how the targeting of cPLA_2_*a* alone or together in treatments with vitamin D analogs affects the system. The model analysis in full is performed and archived as a Jupyter Notebook to ensure reproducibility.

#### 2.2.3 Validating model performance - Stable states & perturbations

With the purpose of validating the performance of the model with respect to its ability to correctly predict experimental observations and its fidelity to biological reality, the stable states or attractors (comparable to cellular states, or phenotypes) of the model under various conditions were computed using the bioLQM toolkit (Naldi, 2018). The cellular states that the model can emulate under various external stimuli can be deduced from the local states of specific subsets of model components that can be used as markers. The balance between the activating and inhibiting states of markers depends on the model’s response to specific inputs and dictates which phenotypes will be reached. For instance, whether a cell will undergo apoptosis does not depend on the activation of pro-apoptotic regulators, but rather on the ratio between pro-apoptotic (e.g. Bax) and anti-apoptotic proteins (e.g. BCL-2) (Khodapasand *et al*, 2015, 2).

The model’s validation and analyses were performed in three main conditions, which are summarized below. In addition to the conditions tested *in vitro* (i.e. the Th17-derived IL-17 and IL-22), a simulation more representative for the psoriatic milieu, where IL-17, IL-22, TNFα, and IFNγ are present, was also included. Additionally, in order to elucidate the potential distinct role of the Th17- and Th1-derived cytokines, the behavior of the model in the presence of the Th1 (i.e., IFNγ and TNFα) and Th17 cytokines separately was explored. Lastly, as EP4 appears to be regulated in response to IL-17 and IL-22 (experimental observations presented in Section 3.1), the receptor was encoded to be active in the simulations.

The three psoriatic conditions that were chosen for the simulations are summarized below:

- **TH17**: Only Th17 cytokines are present, and EP4 is active.
- **TH1**: Only Th1 cytokines are present, and EP4 is active.
- **ALL**: All Th17 and Th1 cytokines are present, and EP4 is active.

As a next step, the behavior of the system under different chemical perturbations was simulated. In all of the three above conditions (i.e. TH17, TH1, ALL), three perturbations were performed:

- **AVX**: Inhibition of cPLA2
- **CAL**: Activation of VDR receptor
- **COMBO**: Combination of AVX and CAL

Regulatory circuits and their functionality were analyzed in order to investigate the existence of multiple attractors and/or cyclic attractors. Functional positive circuits (or *positive feedback loops*) can give rise to multiple attractors, while functional negative circuits (or *negative feedback loops*) can possibly result in cyclic attractors (Thieffry, 2007).

#### 2.2.4 Assessing the system’s state evolution over “time” through stochastic simulations

The probabilistic interpretation of node states, phenotypes, and their potential transient behavior were explored with stochastic simulations using MaBoSS (Stoll *et al*, 2017). Additionally, the ‘time’-dependent evolution of the markers’ states together with the probability of reaching a phenotype were analyzed. MaBoSS allows the calculation of phenotype probabilities when model variables, such as node states or inputs, are altered, allowing the comparison of how the behavior of a system depends on those changes. Stochastic simulations were performed for three conditions; with only Th17 cytokines, with only Th1 cytokines, and with all cytokines present. The analysis was performed to identify whether the evolution of the system could vary depending on the inputs, or whether the system would reach a psoriatic state with either of the two sets of cytokines. In order to ensure that the whole state space was explored, a high number of trajectories (10.000) was chosen. The fraction of instances (from a total of 10.000) a state is occupied is taken as the probability of that state. In a cell population cells may occupy various states with different sets of entities active in each cell, which may result in a different response to the same input, depending on the specific state of that cell. For the majority of the analyses, and unless stated otherwise, the initial state of the cells was not taken into account for the simulations, and the evolution of the system, as well as the probabilities of reaching a state, were calculated while all entities, but the chosen inputs, were inactive.

#### 2.2.5 Understanding the impact of each cytokine in the system through value propagation analysis

A comparison of the impact of Th17- and Th1-derived cytokines was investigated by a value propagation study, as described in (Hernandez *et al*, 2020). The same analysis was also performed for each cytokine individually. The latter comparisons were conducted to identify whether cytokines produced by the same Th-cell populations have complementary or overlapping actions and to compare the influence of each cytokine on its own. In value propagation analysis, a value of 1 is assigned to a node of interest (emulating its activity), in our case the different cytokine receptors of the model, and propagated through the model. This allows the identification of those nodes that are bound to be active or inactive in one of the two compared conditions or in both, and, therefore, aids the comparison of the impact of specific inputs to the system. While setting a value of 1 to the node(s) of interest, the value of the VDR node was set to 0 in all comparisons. This ensured that the analysis would include the influence only of the disease-related nodes, as the effect of Vitamin D analogs had been encoded in the model as an input node that regulates the state of downstream nodes as any other.

#### 2.2.6 Identification of perturbations to restore a normal phenotype

Finally, an analysis was performed in order to explore the complete node perturbation space (both singly and combinatorial) and identify potential perturbations (activating or inhibiting) that would drive the diseased system in a preferred direction and drug targets that could potentially be explored as treatment options. This was performed as a mutation analysis with the Pint tool (Paulevé, 2017), which identifies perturbations that will prevent the activation of the proliferation and inflammatory markers, and the inactivation of apoptotic and differentiation markers, essentially signifying the return to a normal KC phenotype. Pint provided a list of combinations for each of the markers. The combinations were then analyzed and ranked with respect to the frequency that a single or combinatorial perturbation was proposed. It should be noted that the tool might propose combinations that are in fact redundant, meaning that the addition of a second perturbation will have the exact same impact as the individual one. The presented list in the results is curated to include only those perturbations with a distinct impact in the system.

### Data Availability

The model and the notebook containing its full analysis are publicly available at https://github.com/Eirinits/psoKC_model and at the GINsim model repository (http://ginsim.org).

## 3. Results

We set out to develop an *in silico* model of psoriatic KC that could be used to describe the development of cytokine-dependent phenotypes of psoriatic KCs and the therapeutic mechanism of action of cPLA_2_α inhibitors alone and in combination with other anti-inflammatory agents. The construction of the logical model started with describing the wide range of experimental observations regarding the eicosanoid signaling taking place in Th17-cytokine-stimulated KCs that the model should reconcile. Prior knowledge on the system together with experimental observations was integrated into a regulatory network which was then used to simulate the response of KCs to external stimuli and predict their effect on the psoriatic environment with respect to the secretion of ligands and intercellular-acting stimuli. A detailed description of the workflow is provided in the Supplementary information.

### 3.1 Th17 cytokines regulate KC differentiation and prostaglandin E2 release *in vitro*

*In vitro* models of psoriasis can be created by exposing monolayers or 3D skin-equivalent cultures to psoriatic cytokines (Desmet *et al*, 2017). We have previously used the immortalized KC cell line, HaCaT, in 3D skin-equivalent cultures to document the role of cPLA_2_*α* in KC proliferation (Ashcroft *et al*, 2020). These cells express both cPLA_2_*α* and sPLA_2_, and have been used to study the regulation of eicosanoid production by Th1 cytokines in the skin (Sjursen *et al*, 2000; Thommesen *et al*, 1998). HaCaT cells are also known to respond to Th17 cytokines (Woo *et al*, 2017) but the role of cPLA_2_*α* in mediating the effects of Th17 cytokines in KCs is not known.

To investigate the role of cPLA_2_*α* in Th17-dependent signaling in KCs we used air-exposed 3D cultures of HaCaT grown in the presence of a combination of IL-17 and IL-22 (from hereon referred to as Th17 cytokines). Immunohistochemical staining of fixed, paraffin-embedded cultures was used to quantify proliferating and differentiating cells, using Ki67 and cytokeratin (CK) 10 positivity respectively. Eicosanoid release was measured by ELISA. The control cultures were 140-180 μm thick, typically comprising 4-5 cell layers; approximately 20% of the cells were Ki67 positive, consistent with being in a proliferative state, while approximately 30% were CK10 positive, indicating early differentiation. Treatment with Th17 cytokines did not affect the thickness of these cultures but, consistent with several studies (Rabeony *et al*, 2014; Nograles *et al*, 2008; Boniface *et al*, 2005; Pfaff *et al*, 2017), affected differentiation, as evidenced by the loss of CK10 expression. The Th17-treated cultures also had significantly reduced Ki67 positivity indicating lower proliferation, which is inconsistent with the hyperproliferative state of the psoriatic epidermis. Treatment with either AVX001, CAL or COMBO had no effect on the thickness, CK10 or Ki67 positivity of the Th17-treated cultures (**Fig 1A-B**).

**Figure 1.**
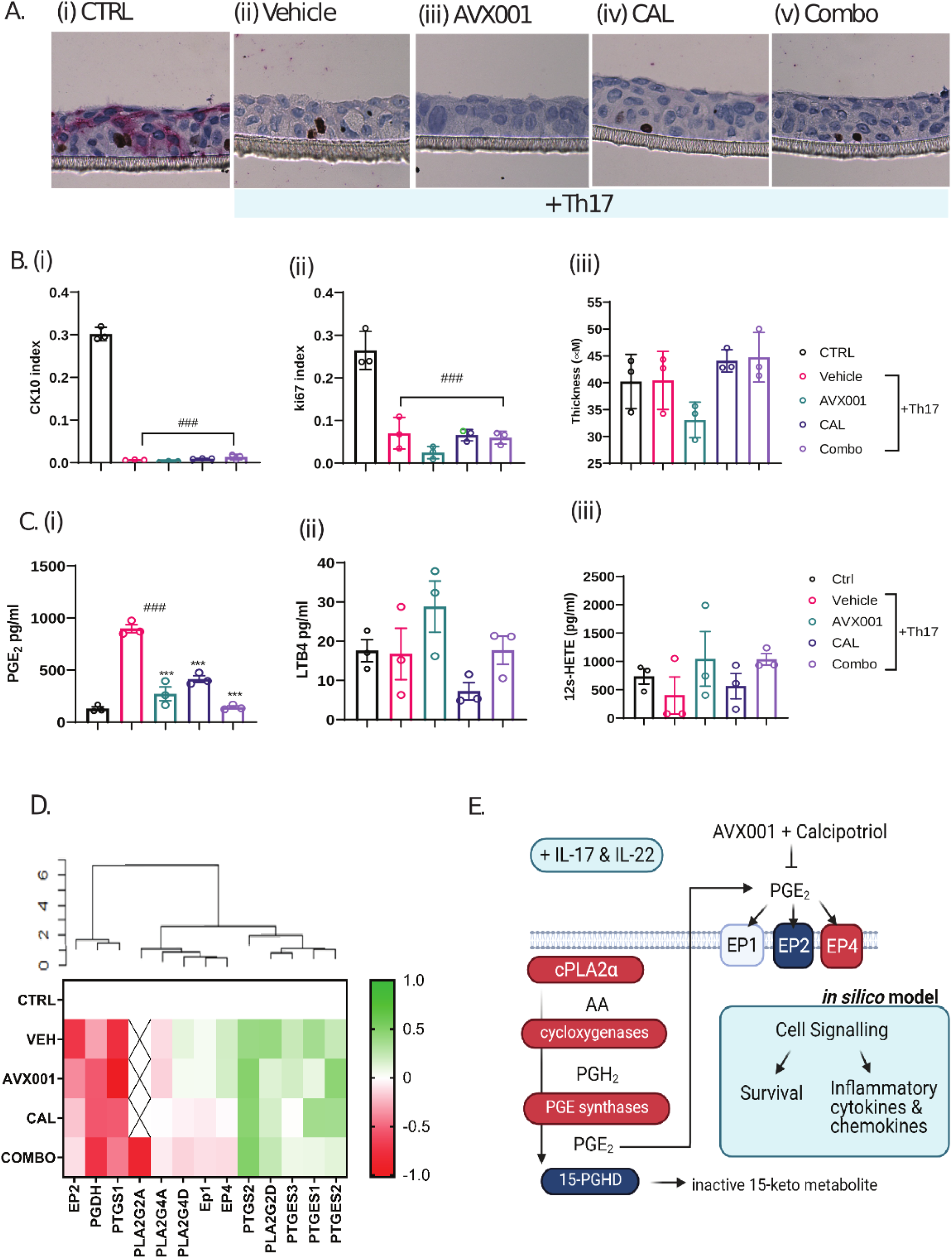
Regulation of cPLA2/PGE/EP4 signaling by Th17 cytokines. A. Representative images showing immunostaining with anti-Ki-67 (DAB+) and anti-cytokeratin 10 (Permanent Red) antibodies B. Quantification of (i) the proportion of CK10 positive cells (ii) the proportion of proliferating cells and (iii) the thickness of the cultures. The data shown are the mean ±SEM for three replicates. C. Eicosanoid levels measured by ELISA for (i) PGE_2_, (ii) 12s-HETE and (iii) LTB_4_. Data shown are the mean ±SEM of three replicates, measured in duplicate. D. Heatmap of Log_10_ fold changes in expression relative to unstimulated controls (CTRL) for genes involved in PGE_2_ synthesis, degradation and signaling. Data is the mean of three replicates. X indicates that expression was undetectable E. Schematic summary showing how Th17 cytokines could regulate PGE_2_ release and activate downstream signaling events via the EP4 receptor. Components in red were up-regulated and components in blue were down-regulated in cultures treated with Th17 cytokines.

Increased levels of the prostaglandin PGE_2_ were measured in the supernatants from cultures treated with Th17 cytokines, while leukotriene B4 (LTB4) and 12s-HETE, were not affected. Suppression of cPLA_2_α activity using the cPLA2α inhibitor AVX001 (Huwiler *et al*, 2012, 00) reduced the PGE_2_ levels implicating the activation of the enzyme in this response. Similarly, the use of the vitamin D analog and topical antipsoriatic drug calcipotriol (10 nM) reduced the PGE_2_ levels, and when we combined both compounds (COMBO) PGE_2_ levels were similar to the unstimulated controls (**Fig 1C**).

### 3.2 Regulation of PGE_2_ biosynthesis and signaling by Th17 cytokines in KCs *in vitro*

The biosynthesis of PGE_2_ results from the co-ordinated regulation of several enzymes; free arachidonic acid (AA) released as a result of phospholipase A_2_ activity is metabolized to PGH_2_ by the activity of cyclooxygenases, Cox-1 (*PTGS1*) and Cox-2 (*PTGS2*), which is further metabolized to PGE_2_ by prostaglandin synthases (*PTGES1, PTGES2, PTGES3*). Intracellular PGE_2_ can be oxidized to an inactive 15-keto form by the enzyme 15-prostaglandin dehydrogenase (PGDH) or released into the extracellular space where it is an autocrine or paracrine ligand for the eicosanoid-prostaglandin (EP) family of receptors (*PTGER1-4)* residing at the cell surface. While, psoriatic lesional skin has been shown to overexpress enzymes involved in PGE_2_ biosynthesis and under-express *15-PGDH* (Lee *et al*, 2019) the separation between regulation of the PGE_2_ biosynthesis pathway in KCs versus infiltrating immune cells was not made.

In order to investigate the mechanism of regulation of PGE_2_ production by Th17 cytokines for integration into the computational model, we measured the expression of genes involved in PGE_2_ biosynthesis and signaling. Total RNA was extracted from the HaCaT cultures treated with Th17, in the absence or presence of AVX001, calcipotriol, or a combination of AVX001 and calcipotriol (COMBO). The expression *of PLA2G4A, PTGS1, PTGS2, PTGES1-3, 15-PGDH,* four prostaglandin E receptors *(PTGER1-4),* and three internal reference genes (*TBP*, *HPRT1*, and *GAPDH*) were measured by quantitative PCR. Results are shown relative to the unstimulated control (CTRL) (**Fig 1D, Fig S2**). Hierarchical clustering of log-transformed values identified 3 groups characterized by upregulation, downregulation, or a lack of regulation by Th17 cytokines. *PTGS2* and *PTGER4* were up-regulated while *PTGS1*, *PTGER2* and *15-PGDH* were down-regulated. A trend towards up-regulation of prostaglandin E synthases was also observed, though no changes reached statistical significance at p<0.05. With the possible exception of *PTGES3*, calcipotriol treatment did not reverse the effects of IL-17 on *PTGS*, *PTGES* or *15-PGDH* expression, indicating suppression of PGE_2_ release by calcipotriol may occur via an independent mechanism. Interestingly, both calcipotriol and COMBO treatments rescued the effect of Th17 cytokines on *PTGER4* and *PTGER2* expression.

In summary, Th17 cytokines stimulated PGE_2_ release from KCs. This was dependent upon cPLA2α activity and was associated with up-regulation of *PTGS2* and down-regulation of *15-PGDH* gene expression. Th17 additionally suppressed *PTGER2* and increased *PTGER4* expression. Combining AVX001 with calcipotriol completely suppressed Th-17-dependent PGE_2_ release and reversed the Th17-dependent switch from *PTGER2* to *PTGER4* expression (summarised in **Fig 1E**). These findings were integrated into the *in silico* keratinocyte model as Th17 cytokine-dependent activation of cPLA_2_*a*/COX-2/EP4 signaling as described below.

### 3.3 The logical model of psoriatic KCs

To recapitulate all these observations in a consistent regulatory model that captures as much as possible the deregulatory events leading to psoriasis we focused on the representation of the KCs as responders to Th cell-derived cytokines. The psoriatic keratinocyte (psoKC) model is presented in **Fig 2**. The model aims to integrate the available knowledge on regulatory interactions which take place during the chronic stages of psoriasis, including the newly described regulation of PGE_2_ signaling. The model contains 90 biological entities (nodes) and 176 regulatory interactions (edges) and can be stimulated by the activation of the receptors recognizing the main psoriatic cytokines, namely IL-17, IL-22, TNFα, and IFNγ, and/or PGE_2_ (EP) receptors. It is important to highlight that the regulatory mechanism of cPLA_2_*a* in the system is mainly designed to be able to assess the effect of PGE_2_ through the EP receptors. This effect is encoded in a way that the activation of EP receptors is an input and not directly activated by the PGE2 node in the model. To account for the autocrine effects of PGE_2_ in KCs, the EP receptors were set to be active in simulations where cPLA_2_*a* activity is uninhibited. In order to distinguish the EP receptor nodes from their genes, which are activated by different transcription factors downstream in the model, a suffix *_g* was added to the nodes that represent their respective genes.

**Figure 2.**
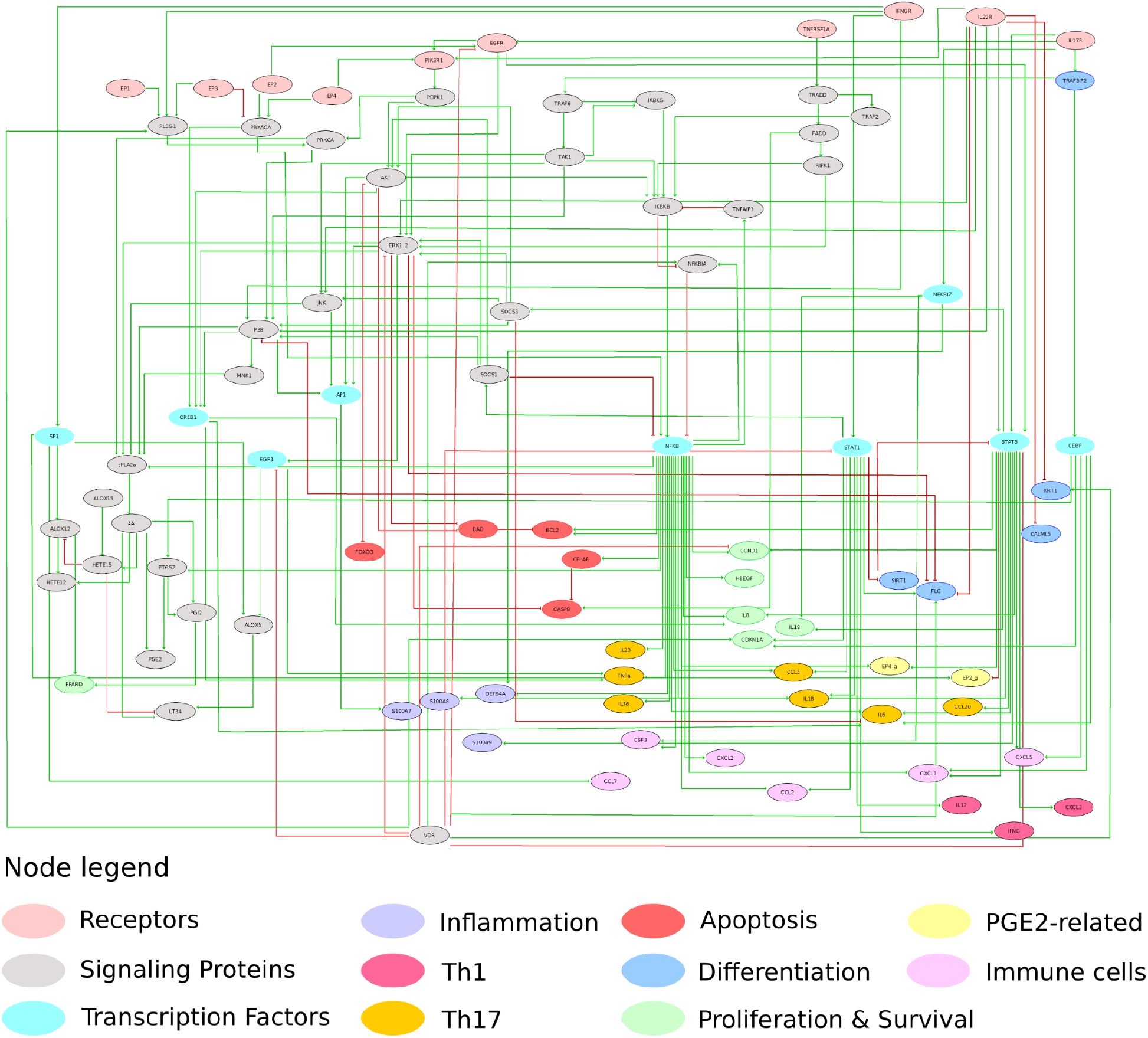
The logical model of psoriatic keratinocytes. The node color depicts their functional role, the phenotype they promote, or the immune cell types they act on. Green lines represent activating interactions and red lines represent inhibitory interactions.

The model is able to describe the three aberrant phenotypes of KC in psoriasis: hyperproliferation, resistance to apoptosis, and aberrant differentiation. The model furthermore covers the KC’s immunostimulatory states, as manifested by the production of cytokines, chemokines, and AMPs (antimicrobial peptides) that activate, recruit and maintain immune cell populations that contribute to the psoriatic phenotype. The phenotypes and state of the system can be inferred from the state (ON or OFF) of selected sets of markers that are characteristic of each specific cellular behavior. The marker nodes were defined and selected as those entities which are included in the transcriptional signatures of the psoriatic cytokines on KCs and associated with one or more phenotypes that the model aimed to represent. The node-cellular state association can be seen in **Table 1**.

**Table 1.** Marker-nodes which were used to define the model’s physiological state and their associated phenotypes and processes. Nodes are named by their HGNC symbol for genes and proteins and their ChEBI ID for eicosanoids. Nodes in red cells are inhibiting the phenotype associated with their respective column. Nodes with an asterisk (*) were experimentally tested.

| Keratinocyte markers |  |  | Inflammatory & immunostimulatory markers |  |  |  |  |
| --- | --- | --- | --- | --- | --- | --- | --- |
| Proliferation & Survival | Differentiation | Apoptosis | Inflammation | Th1 | Th17 | Other immune cells | PGE2-related |
| **CCND1\*** | **FLG\*** | **BAD\*** | S100A7 | **IFNG\*** | **IL-1B\*** | **IL6\*** | **PTGER2_g\*** |
| **CXCL8/IL8\*** | **KRT1\*** | CASP8 | S100A8 | CXCL3 | **TNFA\*** | **CXCL1\*** | **PTGER4_g\*** |
| **PGE2\*** | **CDKN1A\*** | **BCL2\*** | S100A9 | IL12 | **CCL2\*** | CCL7 |  |
| IL19 ** | SIRT1 | **CFLAR** | DEFB4A |  | **CCL5\*** | CXCL2 |  |
| 12-HETE | CALML5 |  |  |  | CCL20 | CXCL5 |  |
| PPARD | **IL36** |  |  |  | IL23 | LTB4 |  |
| HBEGF | **TRAF3IP2** |  |  |  | IL36 | CSF3 |  |
| **CDKN1A** |  |  |  |  |  |  |  |
| **SIRT1** |  |  |  |  |  |  |  |

\*\* IL-19 activates fibroblasts to produce keratinocyte growth factor, which promotes keratinocyte proliferation.

### 3.4 Integration of experimental observations into the model & validation with in vitro results

Our experimental observations suggest that cPLA_2_*a*/PGE_2_/EP4 signaling is active in response to Th17 cytokine in KCs. The potential involvement of this pathway in the development of psoriasis was tested in model simulations by comparing the states of phenotypic marker nodes when EP4 was active and inactive. In conditions where EP4 was inactive, CFLAR, CREB1, IL-8, CSF3, DEFB4A, IL-23, IL-36 genes were also predicted to be inactive. However, upon the activation of EP4 in the simulations, the states of the aforementioned entities were corroborating the available literature on the state of these entities in psoriasis.

In order to further investigate the role of cPLA_2_*a*/PGE_2_/EP4 signaling experimentally, the expression of 17 phenotypic marker genes shown in Table I was measured by qPCR. Th17 up-regulated the expression of pro-inflammatory cytokines (*IL6, IL1β, TNF-α, IL-8, CCL2, CXCL2*) and down-regulated genes associated with differentiation (*FLG, KRT1*) and apoptosis (*BAD*) (**Fig 3** and **Fig S3**). Inhibition of the cPLA_2_*a* using AVX001 or treatment with calcipotriol alone suppressed the induction of *CCL2* and prevented the down-regulation of *BAD*; calcipotriol additionally caused a partial rescue of the loss of *KRT1* expression. Combining AVX001 with calcipotriol inhibited more of the proinflammatory markers (*CCL2, IL-6,* and *IL-8*). To compare these experimental results to the Boolean states of the corresponding nodes in the model, the gene expression data were discretized to 1s and 0s (As described in **Fig S4**). For 59 of the 68 experimental observations, the model predictions were in agreement (**Fig 3**). Interestingly, the nine observed discrepancies were seen only in perturbed simulations, with four of these nine discrepancies occurring with the use of the cPLA_2_*a* inhibitor alone.

**Figure 3.**
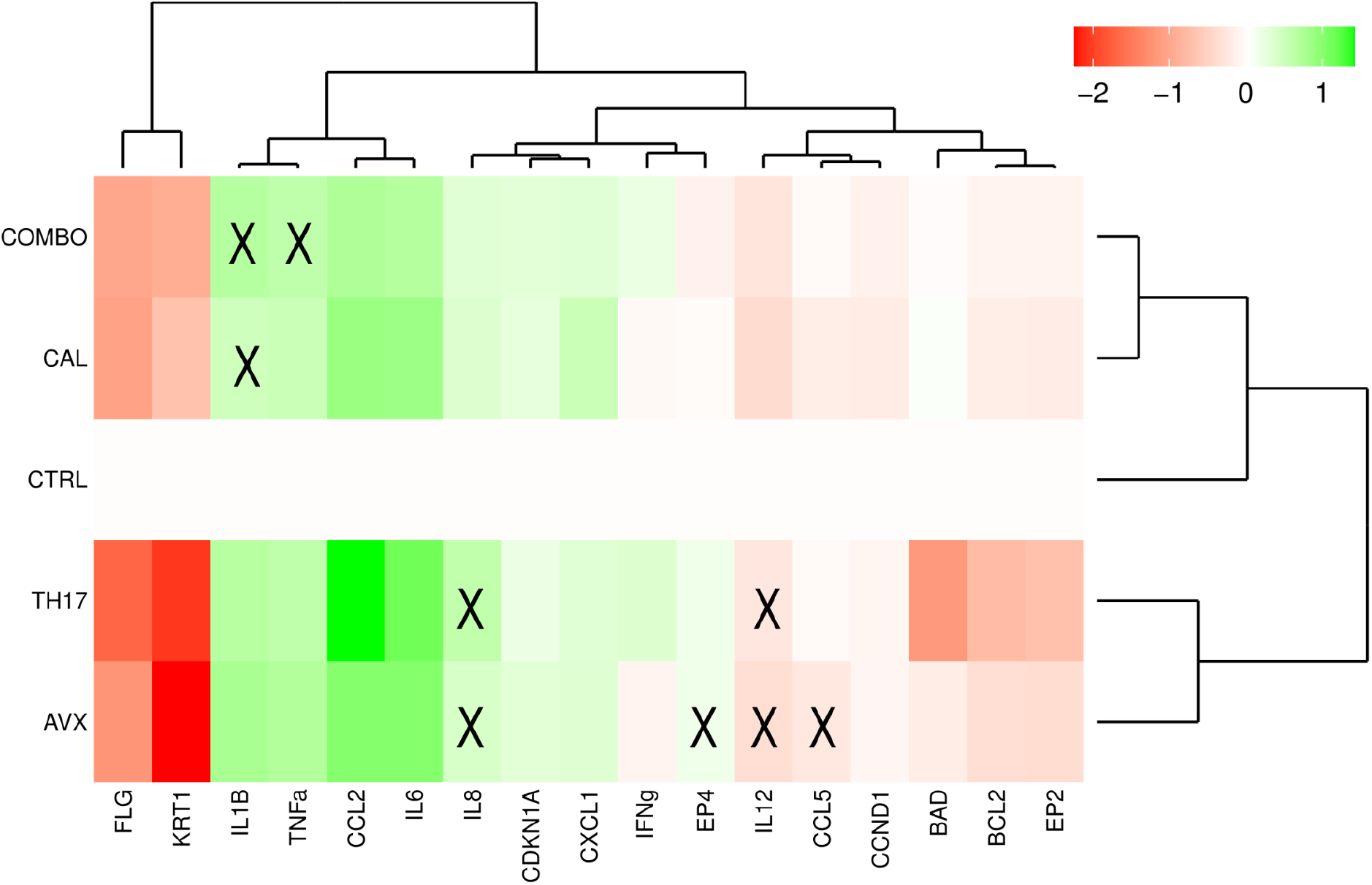
Comparison of experimental observations with node states predicted by the computational model. Heatmap of log_10_ fold change in gene expression relative to CTRL as determined by qPCR. HaCaT 3D cultures were treated with Th17 cytokines (TH17) in the presence of AVX001 (AVX), calcipotriol (CAL), or a combination of AVX001 and calcipotriol (COMBO). Rows and columns are hierarchically clustered. X represents data points where the results from the computational model do not agree with the experimental observation.

### 3.5 Predicting keratinocyte behavior upon different stimuli and treatments

Encouraged by the apparent value that the logical model has in predicting verifiable experimental results, we performed a series of model simulations in an effort to gain further insight into the regulatory mechanisms underlying psoriasis. As previously described, Th17 cells and their produced cytokines play a critical role in psoriasis. However, it has been suggested that different Th-cells types are dominating and controlling the inflammatory process during the various stages of the disease (Furiati *et al*, 2019), but in reality, Th1, Th17, and Th22 cells all together contribute to its development (Diani et al., 2016). Therefore, the downstream simulations and experimentations were focused on gaining a better understanding of how different sets of cytokines, and by extension, different sets of Th cells, affect the behavior of KCs, how this behavior changes when all Th1 and Th17 cytokines are present and, lastly, how PGE_2_-regulated signaling is integrated into the system.

The response of KCs to stimuli from Th1 (i.e. IFNγ and TNFα), Th17 (i.e. IL-17 and IL-22), and Th1/Th17 combined was simulated. For the same conditions, the effect of chemical perturbations with cPLA_2_*a* inhibitors, vitamin D analogs, or a combination of the two was simulated. The simulation results are presented as a series of active and inactive entities for each condition in **Fig 4**. All perturbed-system simulations were able to reach a single stable state, a phenomenon that can be attributed to the absence of positive regulatory circuits (Thieffry, 2007), as identified by a functional circuit analysis performed on the logical model.

**Figure 4.**
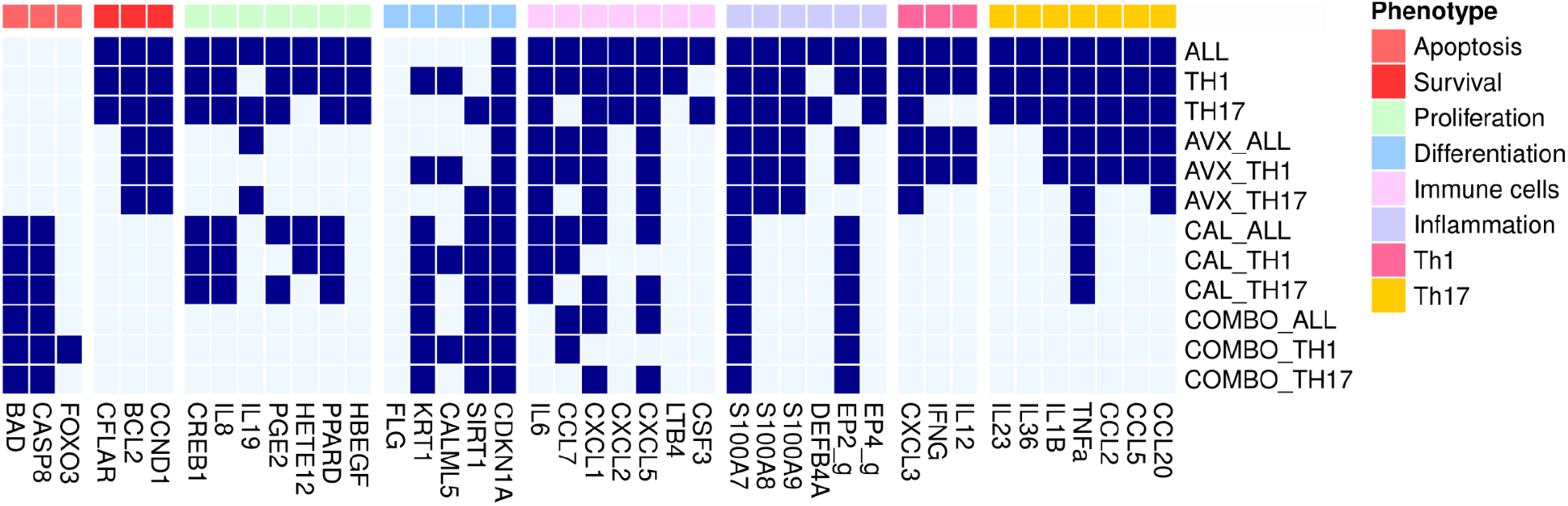
Heatmap of system’s perturbations. The heatmap depicts the set of active and inactive marker nodes, grouped by their associated phenotype, in each simulated condition. Light blue denotes inactive entities while dark blue denotes active entities, the number of which can be taken as a measure of compliance with specific phenotypes.

As displayed in the first three rows of the heatmap, which represent unperturbed conditions, the model predicts that all cytokines contribute to the maintenance and amplification of the positive feedback loop between KCs and Th cells. However, it is only when all four cytokines are present (ALL condition in **Fig 4**) that all immunomodulatory markers are activated. The significant overlap between the Th1 and Th17 conditions could indicate that the two sets of inputs act synergistically rather than complementary, as similar observations were made when analyzing the synergistic effect of IL-17, TNFα, and IFNγ (Chiricozzi *et al*, 2011). Interestingly, the inhibition of cPLA_2_*a* (rows 4-6) mainly affects proliferation markers and only some inflammatory and immune cell markers. These results agree with previous studies of the role of PLA2 enzymes and their downstream products in psoriasis (Ashcroft *et al*, 2020) and skin biology in general (Murakami *et al*, 2018). Moreover, it is worth noting that the targeting of cPLA_2_*a* appears to mainly act on the Th17 cell-related markers, but not on the Th1 markers. However, as the model is mainly focusing on the role of PGE_2_ in the system, it is possible that other cPLA_2_*a* downstream products affect Th1 cell markers, but that their associated mechanisms are not depicted in the current version of the model.

When comparing the effect of cPLA_2_*a* inhibition with the effect of calcipotriol (rows 4 - 6 and 7 - 9 in **Fig 4**), it is evident that both drugs have a distinct mechanism of action, as calcipotriol appears to affect a different set of markers than cPLA_2_*a*. More specifically, calcipotriol affects mostly differentiation and apoptosis, while cPLA_2_*a* appears to mostly affect proliferation. Calcipotriol is able to have both an antiproliferative (Kristl *et al*, 2008; Liang *et al*, 2017) and a pro-apoptotic effect (Huang *et al*, 2019; Tiberio *et al*, 2009) in psoriatic KCs. Apoptosis is subject to an elaborate regulation and controlled by the ratio of pro- to anti-apoptotic regulators, meaning that both types of regulators can be active at the same time (Jan & Chaudhry, 2019). As seen in **Fig 4**, calcipotriol appears to shift the balance towards pro-apoptotic regulators and appears to promote apoptosis in the system, regardless of the input conditions. At the same time, proliferation-promoting eicosanoids, such as PGE_2_ and 12s-HETE, are predicted to still be produced, unless cPLA_2_*a* is inhibited. The same behavior has been reported in various dying cells, where PGE_2_ is released as a Damage-Associated Molecular Pattern (Hangai *et al*, 2016). Keratinocyte differentiation is also rescued by calcipotriol, as indicated by the states of the differentiation markers. Interestingly, terminal differentiation of KCs shares many similarities with apoptotic mechanisms (Terskikh & Vasil’ev, 2005) as can be seen in the coordinated change of both sets of markers.

Finally, as seen in the COMBO simulations, a distinct change of the markers’ behavior is evident, both in the KC cell fate and the suggested effect of the KC cell on immune cell behavior. While we cannot directly quantify the effect of the different drugs and their combination, it is clear that the combination of the two drugs results in what appear to be additive changes in the system. cPLA_2_*a* inhibitor and calcipotriol together inhibit all endogenous and exogenous markers that would promote proliferation and survival. Furthermore, the recruitment of immune cells appears significantly impaired (absence of chemoattractants and immunostimulating cytokines), while primary proinflammatory cytokine TNFα is also inhibited when both drugs are used. The effect on differentiation, however, can be attributed solely to calcipotriol, as the additional cPLA_2_*a* inhibition in the combo treatment has no effect on these phenotype markers.

### 3.6 Evolution of the regulatory system through “time”, and phenotype probabilities

While a stable state analysis provides great insights into which stable states a system can occupy, intuitively analogous with cellular phenotypes, stochastic simulations approximate the observation of transient behaviors and the evolution of the states of the nodes until the model reaches those stable states.

As observed in our *in vitro* experiments and reported in the literature (Ekman *et al*, 2019; Pfaff *et al*, 2017; Boniface *et al*, 2005), stimulation by Th17 cytokines, and mainly IL-22, completely suppresses differentiation and apoptosis in KCs, with survival and anti-differentiation markers being active, and differentiation markers inactive with a 99.9% probability in the stochastic simulations. The promotion of an antiapoptotic phenotype by Th1 cytokines was also confirmed by stochastic simulations, where an anti-apoptotic phenotype was reached with 100% probability (**Fig S5**). Nevertheless, other studies have shown that under certain conditions, TNFα and IFNγ can induce apoptosis in KCs (Viard-Leveugle *et al*, 2013; Reinartz *et al*, 1996). To see if this behavior was dependent on the state of specific components of the system, simulations with active TNFα and IFNγ were performed with an exhaustive series of initial conditions where all entities had a 50% chance of being active. While the system was now able to reach several states with different sets of markers being active each time, the states could be separated into apoptosis and survival. By analyzing the difference between those states, two proteins seemed to be determinants for whether apoptosis would be reached; SOCS1 and SOCS3. Cell populations with active SOCS1 and SOCS3 were able to escape apoptosis (**Fig 5**). Both proteins were found overexpressed in psoriatic skin when compared to normal skin (Federici et al., 2002), and their activity may be contributing to the resistance of psoriatic KCs to cytokine-induced apoptosis, via a mechanism involving the activation of PI3K/AKT and NFκB pathways (Madonna *et al*, 2012). Simulations of the dynamic interplay and temporal evolution of the states pro-apoptotic and anti-apoptotic markers revealed that the combinatorial treatment, which inhibits the survival markers, renders pro-apoptotic marker activity unconstrained so that they reach a final active stable state (Supplementary **Fig S6**).

**Figure 5:**
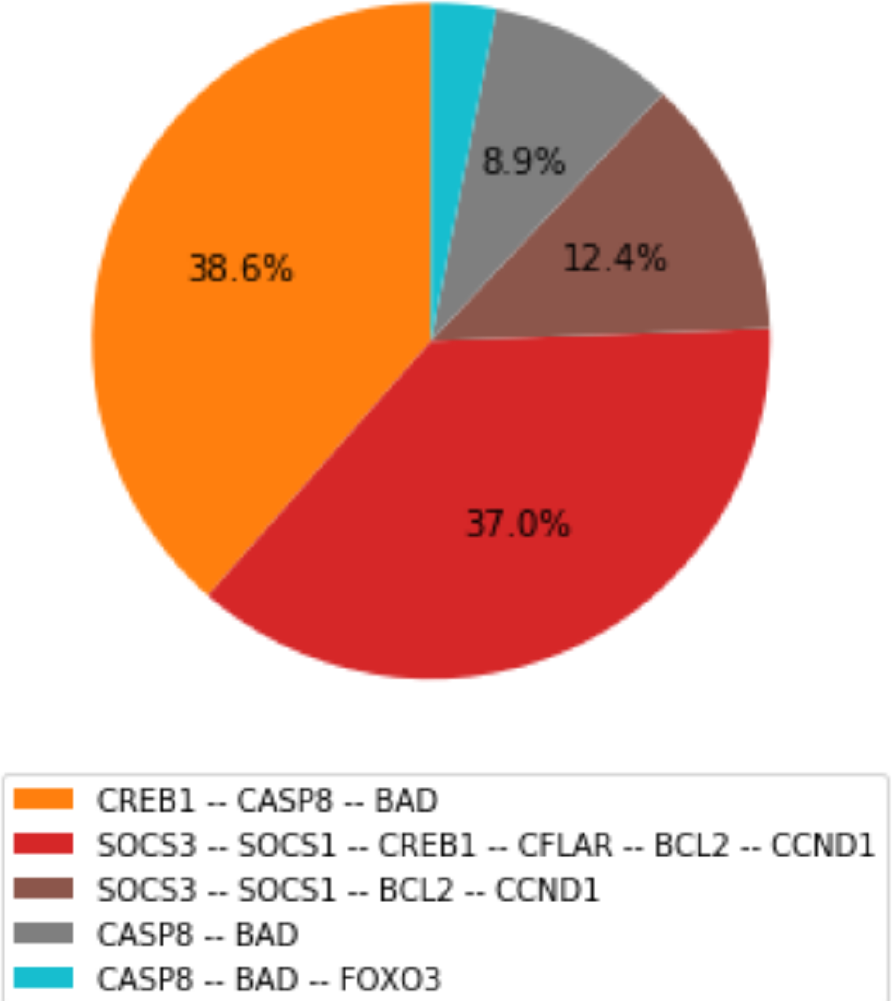
Stable state probability pie chart. Probabilities of stable states after the stimulation with IFNγ and TNFα in an unsynchronized population of cells as calculated by stochastic simulations. Only selected survival and apoptosis markers are shown.

### 3.7 Understanding the impact of each cytokine in the system

To explore in more depth the mechanistic regulation of different inputs that leads to the distinct phenotypes identified by the previous analysis, a ‘value propagation’ analysis was performed allowing to trace in detail the effects of various inputs through the regulatory graph. An analysis of these effects can be used to identify which specific entities are bound to get activated or inactivated by which specific inputs. A comparison of the effects of the inputs should then indicate whether one or more cytokines that derive from the same Th-cell population are sufficient to induce characteristic psoriatic phenotypes or whether it is rather the integration of the stimuli from different Th-cell populations that results in these phenotypes.

The analysis revealed that stimulation with the Th1-derived cytokines impacted more phenotypic markers than stimulation with IL-17 and IL-22 (*see* **Table 2** and **Fig S7** in Supp Mat), as also seen in the stable state analysis. This result agrees with transcriptomic studies that have shown a dominance of Th1 and, more specifically, of IFNγ signature expression profiles in psoriatic lesions (Albanesi *et al*, 2018). However, the transcription of genes of key inflammatory mediators, such as S100 antimicrobial peptides, IL-6, and TNFα, which act synergistically to maintain an inflammatory response in psoriasis, can be activated by both sets of cytokines. The effects of the combination of IL-17 and IL-22 on differentiation and proliferation in the system that was observed in the stable state analysis (*see* **Fig 4**) and *in vitro* experiments (*see* **Fig 1**) are confirmed by the value propagation study, where their simulation primarily fixes the activities of differentiation and proliferation markers to states that are associated with a repressed differentiation phenotype. Conversely, IFNγ and TNFα mainly fix the state of a group of nodes that is associated with Th cell maintenance. At the same time, all cytokines together lock inflammatory nodes in their active state.

**Table 2.** Impact of cytokines in keratinocyte phenotypes and physiological states as identified by a value propagation analysis. The number of nodes whose activity is fixed upon simulations by A) Only IL-17 and IL-22 B) Only TNFα and IFNγ C) All four cytokines. The involvement of the nodes in processes is indicated in the different columns.

| Fixed by.. | All nodes | Proliferation/ Survival | Differentiation | Apoptosis | Th1 | Th17 | Other immune cells | Inflammation | Eicosanoid signaling |
| --- | --- | --- | --- | --- | --- | --- | --- | --- | --- |
| IL-17 & IL-22 | 12 | 1 | 4 | - | - | - | - | - | 3 |
| TNFα & IFNγ | 20 | - | 1 | - | 2 | 3 | 2 | - | 5 |
| IL-17, IL-22, TNFα & IFNγ | 38 | 4 | 1 | 3 | 1 | 3 | 2 | 4 | 5 |

As cPLA_2_*a* is of special interest as a psoriatic drug target, cytokine effects on eicosanoid production were analyzed in more detail. While all cytokines appear to be able to activate the phospholipase, the inducible enzyme COX-2, which catalyzes the production of PGE2, is activated only downstream of IL-17 and IL-22, corroborating our *in vitro* observations. Conversely, TNFα and IFNγ induced the expression of LTB4 and 12-HETE by regulating their respective enzymes. Both LTB_4_ and 12-HETE are reported to play a role in the pathogenesis and development of the disease and have chemotactic properties (Nicolaou, 2013), again confirming the immunostimulatory action of Th1 cytokines. Lastly, 15-HETE, an eicosanoid with anti-inflammatory properties (Nicolaou, 2013), is found to be inactivated in all simulations and value propagations.

A comparison of the distinct effects of IL-17 and IL-22 revealed a greater overall impact of IL-17 on the regulation of both signaling proteins and markers (**Fig S7**), an observation that also finds support in the literature (Nograles *et al*, 2008; Rabeony *et al*, 2014). Nograles et al. also reported that IL-22 but not IL-17 regulated terminal differentiation markers of KCs (i.e. CALML5, KRT1, and FLG). Indeed, upon activation of IL-22, CALML5 and KRT1 are fixed in an inactive state, while proliferation and anti-differentiation markers (i.e IL-6 and IL-29) are active. However, a clear distinction of their role was not observed in our model analysis since both cytokines appear to have an impact on the immunostimulatory markers. In the comparative value propagation between Th1 cytokines (**Fig S8**), it was found that while IFNγ dominates the regulation of immunostimulatory markers, TNFα has a more limited impact.

### 3.8 Model-based analyses to assess possibilities for the treatment of psoriasis

The availability of a logical model representation of psoriatic KCs allows a model analysis that supports an exploration of the possible perturbation space to search for model nodes that may serve as potential drug targets in treatments that could restore a normal phenotype. A full analysis of the perturbation space was performed, with each entity being considered a potential drug target, to allow the identification of novel entities whose targeting should be further explored. The activating or inhibiting, single or combinatorial perturbations that prevent the activation of the proliferation and inflammatory markers, and the inactivation of apoptotic and differentiation markers, are presented in Table 3. The complete lists of perturbations are available in the accompanying Jupyter Notebook.

**Table 3.**
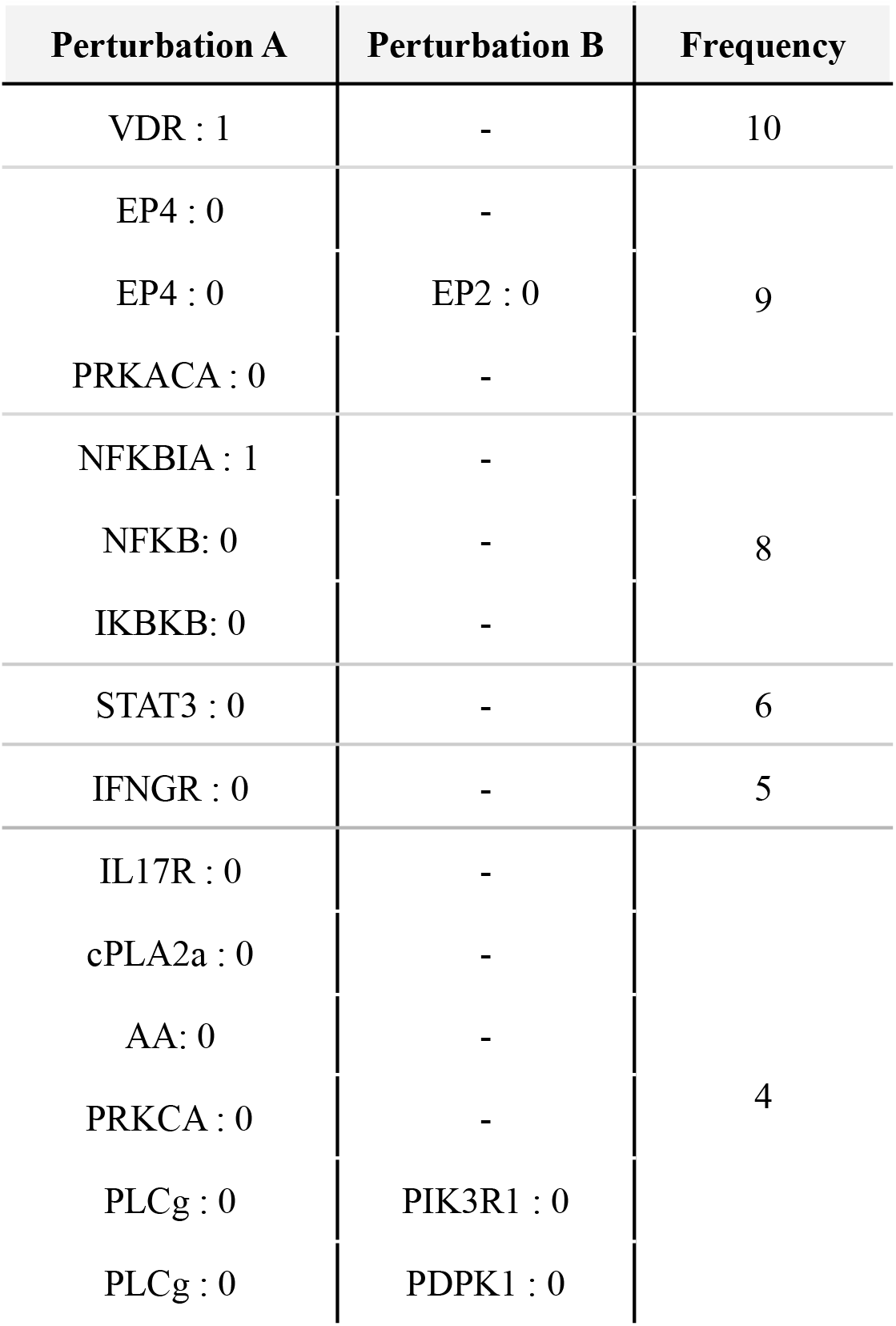
Frequency of perturbations that prevent the system from reaching any of the dysregulated phenotypes in psoriasis. Single or combinatorial, activating or inhibiting perturbations as predicted by PINT.

Generally, the analysis identified the targeting of key inflammatory regulators as the most impactful, especially those that are converging points of multiple pathways and receptors that appear to regulate critical entities for the behavior of the system. Vitamin D analogs such as calcipotriol have been widely used to treat moderate to severe psoriasis, with generally high effectiveness (Kim, 2010). It comes as no surprise, therefore, that the activation of VDR, the receptor activated by calcipotriol, stands out as a key node that can resolve the aberrant phenotypes represented in our model.

A second important perturbation is the inhibition of the EP4 receptor, alone or in combination with EP2, confirming the important role of PGE_2_ and, subsequently, of cPLA_2_*a* inhibition. As endogenous PGE_2_ is not directly activating the EP receptors in these perturbation simulations, but that their activation is fixed in the model when setting the input for the simulations, we suggest that when Pint predicts a perturbation that includes EP receptors, cPLA_2_*a* inhibition could be expected to have the same, or at least similar, effect in the system. The inhibition of cPLA_2_*a* or the inhibition of AA production indeed also appears among the beneficial results, although with a lower frequency. In addition to its proposal as a potential drug target by the model, the direct inhibition of EP4 is increasingly being studied as a potential treatment option in cancer, with its inhibition showing promising results alone or combined with well-established treatments (Yamamoto *et al*, 2020; Konya *et al*, 2013). Based on our model’s predictions, and experimental observations, its inhibition in systems where inflammation is a driver could be a promising option for treatment and should be further tested. The next proposed perturbation was PRKACA (protein kinase cAMP-activated catalytic subunit alpha), with the same frequency as EP4. In the model, PRKACA exerts its effect mainly by regulating key survival pathways and regulators, such as the MAPKs and CREB1. While its involvement in the regulation of proliferative pathways and psoriasis has been documented (Gudjonsson *et al*, 2010), the prediction that it could serve as an additional target that could reduce keratinocyte hyperproliferation in psoriasis is novel and not yet described in the literature.

As NFκB is a convergence point in the inflammatory response and directly regulates the transcription of many inflammatory genes, its direct inhibition or indirect inhibition through its regulators is no surprise as this would restore a more normal phenotype in KCs. The importance of NFκB as a key transcription factor in chronic inflammatory diseases, including psoriasis (Goldminz *et al*, 2013) has made NFκB and its regulators an attractive target for treatment in many diseases (Gilmore & Herscovitch, 2006). Its inhibition, in combination with the inhibition of other transcription factors such as STAT3, has also been explored (Andrés *et al*, 2013). While the combination of NFκB and STAT3 is not detected as an impactful perturbation, the inhibition of STAT3 singly has the next highest frequency in the list. The model proposes the inhibition of STAT3 as well as the inhibition of other entities that control many inflammatory genes that sustain the inflammatory vicious cycle of psoriasis and have been previously studied as potential drug targets. These entities also include the abundantly-studied IL-17 (Amin *et al*, 2018). Indeed, several biologics used in psoriasis are targeting IL-17 (Ly *et al*, 2019). At the same time, while IFNγ exerts a significant influence on the system, it has been reported that its inactivation alone is not enough to convert the psoriatic phenotypes back to normal, while its inhibition together with other entities such as IL-17 appears to be more effective (Meephansan *et al*, 2017). At the bottom of the list, some perturbations suggest the targeting of PI3K/AKT pathway components together with PLCγ, both pathways linked to proliferation and cell survival (Castilho *et al*, 2013; Haase *et al*, 1997).

## 4. Discussion

In this study, we aimed to develop an executable logical model for investigating the regulation of keratinocyte physiology by pro-psoriatic cytokines that could be used to investigate the therapeutic mode of action of cPLA_2_α inhibitors in psoriasis and predict likely druggable combinatorial partners for future investigations. Our model, supported by primary experimental observations, suggests that cPLA_2_α-dependent PGE_2_/EP4 signaling is important for maintaining the psoriatic phenotype of KCs under cytokine stimulation, thus providing a therapeutic mode of action for the cPLA_2_α inhibitor AVX001 in psoriasis. Furthermore, we suggest AVX001 and the topical antipsoriatic drug calcipotriol have overlapping and distinct mechanisms of disease resolution and predict beneficial therapeutic effects when used in combination.

The pathology of psoriasis involves an interaction between immune cells and epidermal KCs that is critical for maintaining the chronic disease state, with keratinocyte activation leading to the release of chemokines and cytokines that promote the infiltration and amplification of immune cells (Lowes et al.). In our analysis we have focused on the key players that drive a vicious cycle of inflammation: the eicosanoids produced by KCs and act on KC themselves or immune cells, Th1-derived cytokines, which are established activators of phospholipases and cause eicosanoid release from KCs (Sjursen *et al*, 2000; Thommesen *et al*, 1998), and Th17-derived cytokines, which have largely unknown effects on eicosanoid signaling.

Our experimental findings demonstrated the importance of Th17 regulated PGE_2_ release via cPLA_2_*a* in the induction of proinflammatory cytokine expression in KCs, and suggest a role in cell survival. The dependence of IL-17 responses on PGE_2_ signaling was previously shown in normal human epidermal KCs and demonstrated to involve the activation of the MAPK pathway (Kanda *et al*, 2004, 2005). Since cPLA_2_*a* is a well-known target of the MAPK pathway (reviewed in (Clark *et al*, 1995)), this presents a putative mechanism for its regulation by Th17 cytokines. Th17 cytokines also upregulate *PTGER4*, which in turn can modulate cell survival and proliferation via PI3K (Peng *et al*, 2017) and ERK (Fujino *et al*, 2003) signaling. This suggests that Th17 cytokines may also regulate how the KCs respond to PGE_2_ and supports evidence that signaling via EP4 may predominate in psoriasis (Lee *et al*, 2019). Interestingly, calcipotriol suppressed the Th17-dependent release of PGE_2_, but not the up-regulation of *PTGS2*, and counteracted the changes in *PTGER4* and *PTGER2* expression. The suppressive effects of calcipotriol on PGE_2_ release differs from previous studies showing that calcipotriol can stimulate, or augment PGE_2_ release in both KCs and immune cells (Ashcroft *et al*, 2020; Ravid *et al*, 2016) and is rather consistent with the inhibition of Th17-dependent pro-inflammatory responses as described by (Lovato *et al*, 2016). It will be interesting to investigate the mechanism of suppression of Th17-dependent PGE_2_ release by VDR activation, given it may be a novel mechanism accounting for some of the anti-psoriatic effects of the compound. The experimental data thus suggest both the cPLA_2_*a*-COX2-PGE_2_ synthesis pathway and PGE_2_/EP4 signaling pathways are active in psoriatic KCs, with implications for both paracrine and autocrine signaling. The impact of autocrine EP4 signaling was further investigated using the psoKC model. Simulations revealed that the states of key inflammatory and proliferation markers were active, corroborating their reported state in psoriasis only in simulations where EP4 was activated. We thus hypothesized that activation of the PGE_2_/EP4 axis is involved in an intrinsic amplification loop that could enable or sustain the effects of psoriatic cytokines and that suppression of the PGE_2_/EP4 axis by inhibition of cPLA_2_α presents a putative mode of action for AVX001 in psoriasis. We further explored this hypothesis using the *in vitro* model system by analyzing the expression of a subset of the phenotypic marker genes assigned as outputs, and by exploring the behavior of the KCs in different conditions using our computation model.

Phenotypic characterization of the *in vitro* model showed its inability to recapitulate the hyperproliferative state of KCs in response to Th17 stimulation, which is a prominent feature of psoriatic skin. Evidence for the usefulness of such epidermis-equivalent cultures to recapitulate the hyperproliferative state of KCs in response to cytokine stimulation is lacking, indicating a potential requirement of either the dermal compartment or alternative immunological signaling molecules to recreate this aspect of the disease (reviewed in Desmet *et al*, 2017). The computational model, on the other hand, accurately predicted the hyperproliferative state of the KCs in response to Th17 cytokines, allowing us to investigate the potential implication of inhibiting the cPLA_2_*a*/PGE_2_/EP4 axis for a wider range of psoriatic phenotypes. Furthermore, the psoKC model was able to accurately reproduce the majority of our *in vitro* observations, in addition to those reported in the literature. We further explored its predictive use by expanding the simulations of the system’s response to additional stimuli that were not tested in the lab, namely IFNγ and TNFα. The ability to capture a more representative cytokine microenvironment for psoriatic KCs, along with the integration of the cPLA_2_*a*/PGE_2_/EP4 axis gave us access to an in silico experimentation system that allowed us to test a wide range of stimuli, observe the propagation of signals through the system and predict the effects of mutations.

Logical models have been previously used to elucidate drugs’ mechanisms of action and effectiveness (Béal *et al*, 2021; Traynard *et al*, 2017). *In silico* treatment with the drugs tested *in vitro* revealed distinct mechanisms of action and interactions between the cPLA_2_*a* inhibition and Vitamin D analogs, where the two drugs impacted different phenotypic aspects of KCs. Calcipotriol is able to have both an antiproliferative (Kristl *et al*, 2008; Liang *et al*, 2017) and a pro-apoptotic effect (Huang *et al*, 2019; Tiberio *et al*, 2009) in psoriatic KCs. The model predicted that calcipotriol, alone or in combination with cPLA_2_*a* inhibition, acts either through rescuing the differentiation phenotype and/or via the induction of apoptosis. Alternatively, the inhibition of cPLA_2_*a* signaling directly impacts the proliferative phenotype of the simulated psoriatic KCs, as already described by (Ashcroft *et al*, 2020). Additionally, as initially hinted by the *in vitro* experiments, the combination of the two treatments uncovered a common regulation of several system components related to both KC physiological state and inflammatory immune response.

While an extensive literature search was employed in order to refine the model in a way that it correctly predicts the experimental observations, at the molecular level, some observed discrepancies between observations and predictions could be attributed to various reasons. First, the discretization method of the experimental results together with the intrinsic abstraction of discrete models could lead to some loss of information about the state of an entity. For instance, we observed experimentally that calcipotriol treatment reduced the expression of Th-17- stimulated *IL1B* expression but the effect was not significant and therefore did not meet the threshold set during the discretization process for inhibition. At the same time, the model predicts that IL-1β will be completely inhibited by calcipotriol. This discrepancy could be explained by the inability of the logical model to distinguish between partial and full inhibition, or that the threshold for discretization based on the experimental observations was too stringent. Additionally, as a gatekeeper of inflammation, IL-1β is under a strict regulation involving post-translational cleavage of proIL-1β, the interleukin 1β protein precursor, for complete activation (Liu *et al*, 2016). The control of calcipotriol over this mechanism of activation has been described in other systems, such as hematopoietic stem cells (Wang *et al*, 2020) and is thus included in the PKN, but not captured by measuring gene expression experimentally. Similar reasoning could explain the wrongly predicted state for IL-8 after cPLA_2_*a* inhibition; PGE_2_ has been reported to regulate IL-8 gene transcription through epigenetic mechanisms in human astrocytoma (Venza *et al*, 2012), and a similar regulation could occur also in KCs. While there was an extensive effort to incorporate the regulatory interactions relevant to eicosanoid signaling reported in the literature, half of the discrepancies involve the state of markers upon treatment with cPLA_2_*a* inhibitors. In addition to the aforementioned discretization limitations and post-translational regulation issues, wrong predictions may also result from the focus of the model on PGE_2_ signaling and not signaling mediated by other eicosanoids, or from gaps in the current knowledge on how cPLA_2_*a*-derived lipid signaling mediators influence KCs.

We further demonstrated how a logical model can be used to better understand a system and its internal regulatory mechanisms. As proposed by Thieffry et al., the presence of negative circuits (i.e. negative feedback loops) in a model is expected to generate cyclic attractors (Thieffry, 2007), where the state of the nodes is oscillating (i.e. continuously alternation between active and inactive states). Remarkably, the model contained a very limited number of functional circuits. This attribute became apparent during the integration of psoriasis-specific information in the model. A characteristic example is the negative feedback loop between STATs and SOCSs, where STAT1 and STAT3 activate their inhibitors SOCS1 and SOCS3. However, the activation of SOCS1 and SOCS3 appears inadequate to completely inhibit the expression of STAT-downstream targets in psoriasis (Madonna *et al*, 2010), making this negative circuit non-functional in the context of psoriasis, and it was, therefore, removed from the model. This observation on the model’s behavior proposes the deregulation of mechanisms that would otherwise limit the spread of inflammatory response that might contribute to the development of psoriasis. Furthermore, stochastic simulations together with value propagation analyses confirmed the distinct role of Th1 and Th17 cells in the pathophysiology of psoriasis. All results indicated that Th1-derived cytokines have a key role in stimulating and further enhancing immune responses by regulating the majority of immune markers and chemotactic eicosanoids, as described in (Albanesi *et al*, 2018). The activation of cytokines and chemokines related to the recruitment, survival, and maintenance of Th17 and Th22 subpopulations by Th1 cells may indicate that Th1 activation precedes the activation of Th17 and Th22 during the development of psoriasis or that Th1 activation functions to amplify Th17 and neutrophil responses, as proposed in (Kryczek *et al*, 2008). The value propagation results also corroborated the dominance of an IFNγ gene signature, where IFNγ dominated the regulation of markers and TNFα had a more limited impact. This observation supports the claims that TNFα is a potentiator and amplifier of IFNγ effects (Albanesi *et al*, 2018). The synergistic effect of TNFα with IL-17 has also been described (Chiricozzi *et al*, 2011). These integrative responses could explain why the influence of TNFα on other nodes is overlapping on regulations from other inputs. However, it is worth noting that TNFα plays an important role in the development of the disease and its targeting remains a promising therapeutic option, but mainly due to its effect on Th17 cells (Furiati *et al*, 2019; Yost & Gudjonsson, 2009).

Predictive logical models have been previously used to propose treatment strategies that could affect the system in a desired way. The suggested perturbations included both known and novel targets, some of them already involved in well-established psoriasis treatments. For the rest of the targets, inhibitory agents are available, and sometimes even approved as drugs, possibly opening attractive opportunities for drug repurposing. The fact that many of the suggested perturbations concern already used or explored targets confirms that the model sufficiently integrates the role of those entities in the system and is able to recognize the importance of certain stimuli and pathways to the progression of the disease. For example, VDR activation by the use of Vitamin D analogs and the blocking of IL-17 signaling are widely used in the treatment of psoriasis (Kim, 2010; Ly *et al*, 2019, 17). The importance of the cPLA_2_*a*/PGE_2_/EP4 axis was further highlighted by the proposal of components of the pathway as a way to restore a normal phenotype. The role of EP4 was previously speculated to be involved in psoriasis and KCs but is not fully explored for its potentials for treating the disease. Therefore, we further propose the exploration of the role of EP4 in psoriasis and its potential as a drug target. Another entity, the catalytic subunit of Protein Kinase A (PRKACA), was proposed as a promising drug target. PRKACA’s downstream effects involve multiple pathways, including WNT signaling, a pathway directly related with cPLA2*a* signaling (Xu *et al*, 2019), and psoriasis in general (Gudjonsson *et al*, 2010). PRKACA has also been studied in the context of cancer, where its aberrant regulation and overexpression have been related to inflammatory activation of Caspase 1, oncogene activation, and elevated PGE2 levels (Almeida *et al*, 2011). In addition to its role in hyperproliferation and inflammation, PRKACA has been associated with drug resistance in breast cancer by supporting the restoration of anti-apoptotic phenotypes of cancer cells (Moody *et al*, 2015). The current knowledge on the activity of the kinase, together with the model’s predictions on the effects of its inhibition in psoriasis, suggest that its exact role in the development of the disease, together with its potential importance as a prognostic marker and/or drug target should be further explored.

While the model proposes certain perturbations, the need for further assessment of the results based on current knowledge and their actual experimental testing remains. For instance, the targeting of PLCγ together with components of the PI3K/AKT pathway is proposed by the model. This comes as no surprise, as their action affects proliferation and cell survival. However, PLCγ has been described as ‘undruggable’ in the literature (Lattanzio *et al*, 2013), as the development of small molecule inhibitors against it appears troublesome. The blocking of IFNγ was also among the proposed perturbations. Even though its importance on the progression of the disease is supported by both the model and prior knowledge, targeting of IFNγ on its own fails to restore a normal phenotype (Meephansan *et al*, 2017). Despite the model providing some insights on intracellular communication by integrating the effects of immune cell-derived cytokines as stimuli and KC-derived secreted factors that influence immune cells, it likely covers only a fraction of the complex intra- and inter-cellular signaling that underlies psoriasis. It is, therefore, expected that some observations at the tissue level cannot be captured to their totality by single-cell models. While understanding the role and behavior of individual cells in the disease, multicellular models are likely to provide a more accurate representation of biology as they would capture emergent behaviors that arise from intercellular interactions and communications.

In conclusion, this paper demonstrates how a combination of *in vitro* and *in silico* models, both without doubt flawed in their accuracy of representing the system to be analyzed, can complement each other to test and generate hypotheses for characterizing the regulatory mechanisms and effects of stimuli in a cellular system. Use of the *in silico* model allowed us to interpret *in vitro* observations in a more holistic manner providing additional mechanistic details of drug actions, and led us to propose novel candidates of drug targets that should be further explored. Through this collaboration, we characterized the regulation of certain lipid mediators in psoriatic KC, which together with the prior knowledge of the involvement of cPLA_2_*a* signaling revealed yet another layer of involvement of KCs in the chronic inflammatory loop of psoriasis. Additionally, we presented a computational framework to interpret the effect of chemical perturbations and various stimuli in a cellular network by exploring the differences between the mechanisms of actions of cPLA_2_*a* inhibitors and calcipotriol. A perturbation analysis revealed promising entities whose targeting could restore a normal phenotype in KCs. For many of these targets, inhibitory agents have already been described, and sometimes these are even available as approved drugs, possibly opening attractive opportunities for drug repurposing. The combination of model-based analyses showcases how a systems biology approach can support a better understanding of a disease, further drive hypotheses on its causes, and suggest new targets for potential treatment.

## Author contributions

ET and MK designed the in silico approach. FA and BJ designed the in vitro approach. ET performed the in silico model development and analysis. FA designed and performed the in vitro experiments. ET and FA designed the model validation experiments. ET and FA wrote the manuscript with input from all authors.

## Acknowledgments

The authors would like to acknowledge Denis Thieffry and Laurence Calzone for their support on the building and analysis of the logical model.

